# The contemporary Pacific gray whale (*Eschrichtius robustus*) gene pool includes ancestry from a potential ghost population: inferences from population genomics

**DOI:** 10.64898/2026.02.17.706390

**Authors:** Natalie M. Allen, Anna Brüniche-Olsen, Jong Yoon Jeon, Andrew J. Mularo, John W. Bickham, Jorge Urbán Ramírez, J. Andrew DeWoody

## Abstract

Local extirpations and extreme bottlenecks can deplete genetic variation and obscure population dynamics, especially in highly mobile species. The gray whale (*Eschrichtius robustus*) undertakes some of the longest known migrations among mammals, and conventional wisdom acknowledges two stocks (eastern and western gray whales) in the Pacific. Commercial whaling depleted both stocks, and while the Eastern North Pacific (ENP) stock has rebounded, the western stock was feared extirpated. In the post-whaling era, the origin of a small summer aggregation near Sakhalin Island, Russia (which we refer to as the Western North Pacific (WNP) stock) is unclear; this group may include descendants from the original western stock, founders from the eastern stock, or some combination of the two. To clarify the genetic affinities of WNP gray whales, we generated whole genome resequencing data for 74 individuals sampled from both geographic regions to assess their origins. Surprisingly, WNP whales are more genetically varied than ENP whales according to principal components and admixture analyses. We present evidence that this structure is due to mixed ancestry in WNP, with admixture occurring between historical eastern and western stocks. Genomic signals based on both single nucleotide polymorphisms and on copy number variants indicate that despite mixed ancestry, the influx of eastern gene flow has largely homogenized genomic diversity across the Pacific. These findings highlight the ability of whole-genome data to help resolve questions of extirpation and to clarify complex gene flow dynamics in highly mobile species.

## Introduction

### Background

Population extirpations, especially in species of conservation concern, can lead to the irreversible loss of genetic variation at both the local and metapopulation scale (McCauley et al., 1991). When a subpopulation is presumed extinct then seemingly reappears, it can be difficult to determine whether the new population represents a recolonization or cryptic persistence, especially in the absence of historical samples. In some cases, extirpated populations may be detectible only in the form of introgression from an unsampled “ghost population”, whereby cryptic persistence or admixture with recolonizing lineages leave behind genetic signatures (Ottenburghs, 2020). Such demographic uncertainty makes it challenging to understand population dynamics, thereby limiting evolutionary insights (including those relevant to conservation efforts). This challenge is further compounded in highly mobile species such as migratory animals, where patterns of gene flow are harder to decipher and movement dynamics often manifest in unexpected ways (Pearse et al., 2019). Genetic evidence in humpback whales (*Megaptera novaeangliae*), for instance, suggests that travel to breeding grounds facilitates gene flow between stocks from different hemispheres (Rizzo and Schulte, 2009). Another study in blue whales recently revealed long-distance connectivity that challenges subspecies classifications (Attard et al., 2024). These examples underscore how vagility can obscure population structure, emphasizing the need for robust datasets to inform conservation efforts and to interpret evolutionary history and taxonomy. Pacific gray whales (*Eschrichtius robustus*) are a compelling case study for this problem given their long-distance movements, historical bottlenecks, and uncertain persistence.

Gray whales were once present in both the Pacific and Atlantic oceans but were extirpated from the Atlantic in the 1700s, likely due at least in part to commercial whaling (Mead and Mitchell, 1984). In the Pacific, two stocks were historically recognized: eastern gray whales, which inhabited the coast of North America with wintering grounds off of Baja California, Mexico, and western gray whales, whose range extended from the Sea of Okhostok to the coastal waters of South Korea (Fig. 1). The western stock was long thought to be extirpated, but a population of whales now exists near Sakhalin Island off the Asian coast. In the western Pacific, gray whales reportedly “disappeared” for more than 35 years (from ∼1933–1970; Weller and Brownell, 2012; Fig. 2b). This prolonged period of nondetection raises the possibility that the contemporary Sakhalin Island whales represent a) remnant (and/or descendant) individuals from the historical western stock; b) long-distance migrants from the eastern stock; or c) admixture between a western ghost population and migrants from the eastern Pacific (thus potentially constituting an evolutionary amalgamation of genetic material from a remnant western ghost population and migrants from the eastern Pacific).

**Figure 1:**
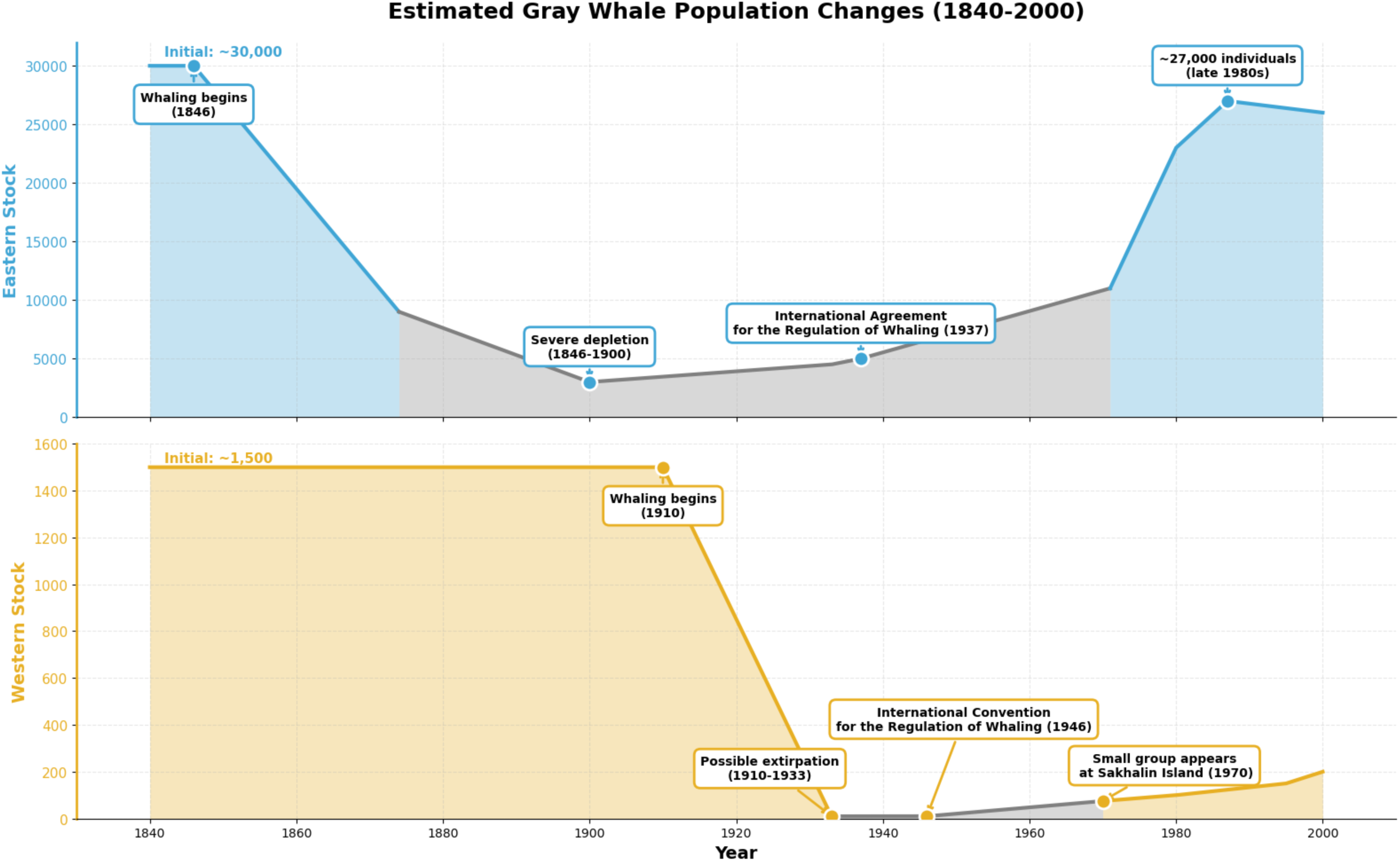
*Conventionally accepted historical range map for eastern gray whales (blue) and western gray whales (yellow) generated with IUCN range data (IUCN, 2024). Overlap in the range is shown in striped yellow and blue. The green shading denotes Sakhalin Island, Russia, and red dots denote sample sites for this study. Gray whale photo taken by Anna Brüniche-Olsen.* Alt Text: Map showing the conventionally accepted historical ranges of eastern North Pacific (ENP) gray whales in blue and western North Pacific (WNP) gray whales in yellow. The ENP range extends from Baja California, Mexico, along the west coasts of the United States and Canada, through the Bering Sea and down the coast of Russia, ending along the east coast of Kamchatka. The WNP range begins along the east coast of Kamchatka, overlapping the ENP range, and extending down along the coastlines of Russia, Japan, Korea, and China to end in the South China Sea. Two red dots denote sample sites off the coasts of Sakhalin Island, Russia, and Baja California, Mexico. The ENP wintering grounds are indicated off the coast of Baja California, Mexico while the presumed WNP wintering grounds are indicated in the South China Sea. A gray whale photograph can be seen in the center of the Pacific Ocean.

**Figure 2:**
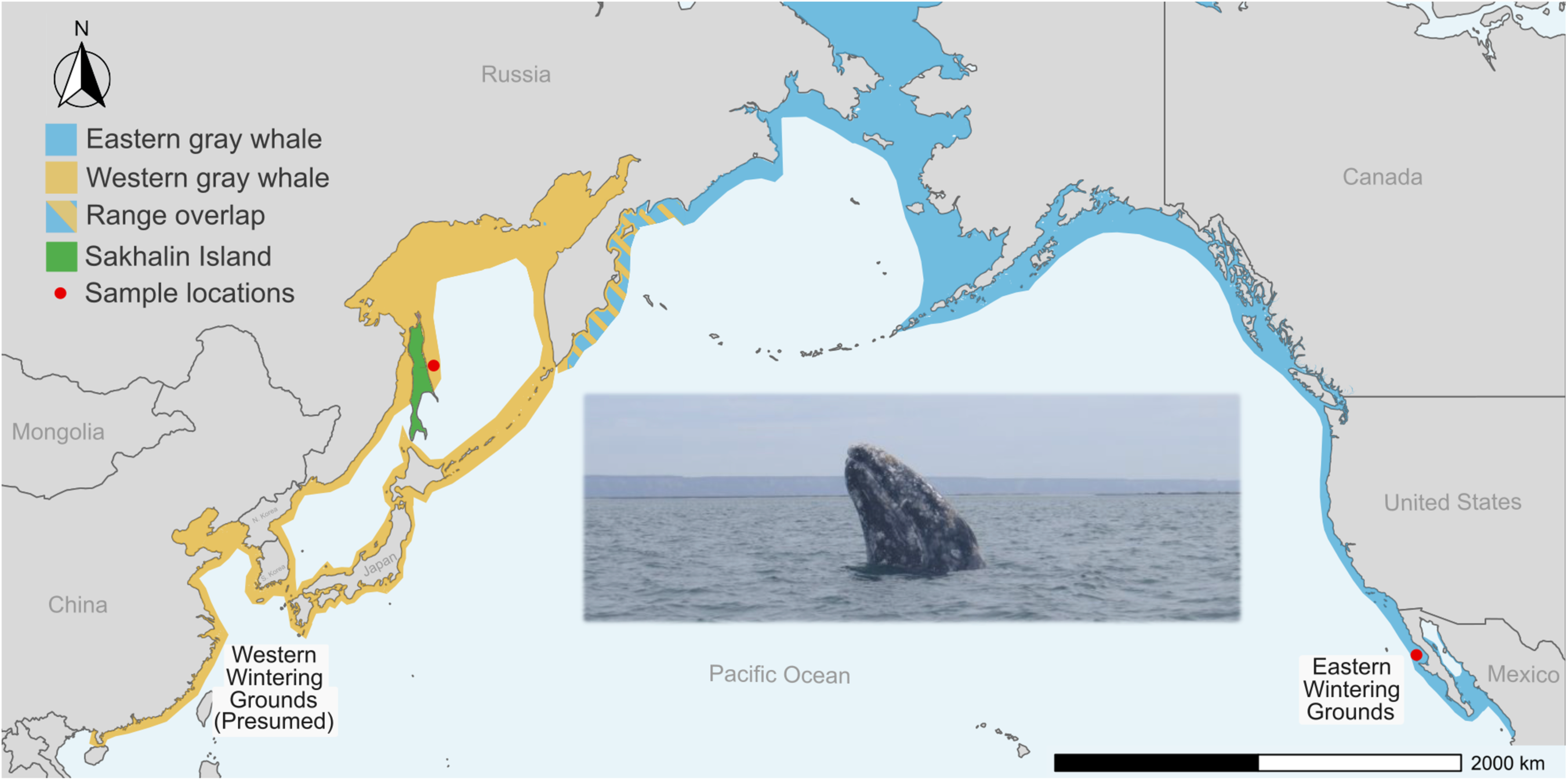
*Approximate census population size changes in the eastern North Pacific (blue) and western North Pacific (yellow), 1840-2000. Gray shading indicates time periods where population estimates are unavailable, and the plot represents a “best guess” at overall trends. The 1937 International Agreement for the Regulation of Whaling ended hunting of gray whales for many countries, but not the Soviet Union or Japan, which ceased whaling after the 1946 International Convention for the Regulation of Whaling. Census estimates were compiled from Rice and Wolman (1971), Scammon (1874), Bowen (1974), and Eguchi (2023), and should be interpreted with caution due to the incomplete nature of population size records. Note that the y-axes have different scales.* Alt text: Line graph showing approximate census population size trends for ENP (blue, above) and WNP (yellow, below) gray whales from 1840 to 2000. The eastern population begins near 30,000 individuals in 1840, declines steeply during commercial whaling to a minimum of ∼3,000 by 1900, then gradually recovers after international protections, reaching ∼11,000 in 1971 and ∼27,000 by 2000; the depletion period (1874–1971) is shaded gray, while pre-depletion and recovery periods are shaded blue, with annotated events marking the onset of whaling (1846), severe depletion, the 1937 International Agreement for the Regulation of Whaling, and late-1980s abundance. The western population begins near 1,500 individuals in 1840, declines to ∼10 by 1933 and remains extremely low through 1946, then increases modestly after 1970 to ∼200 by 2000; the near-extirpation period (1933–1970) is shaded gray, with pre-decline and recovery periods shaded yellow, and annotations marking whaling onset (1910), possible extirpation, the 1946 International Convention for the Regulation of Whaling, and the appearance of a small group at Sakhalin Island in 1970.

Because the ancestry and stock affiliation of contemporary whales are uncertain, we distinguish historical (pre-whaling) eastern and western stocks from present-day whales. Hereafter, we use the terms Eastern North Pacific (ENP) and Western North Pacific (WNP) as geographic descriptions for contemporary whales sampled in the eastern Pacific and near Sakhalin Island, respectively. The ambiguity in population structure underscores the complexity of gray whale conservation efforts. Under current assessments, whales in the WNP are listed as Endangered western gray whales by the IUCN, whereas eastern gray whales are listed by IUCN as Least Concern (IUCN, 2024; IUCN, 2017).

#### Box 1.

##### History of gray whale populations

In the Pacific, western and eastern gray whales have been historically recognized as separate stocks by organizations including the International Whaling Commission (IWC) and the International Union for the Conservation of Nature (IUCN) (Rice and Wolman, 1971; Fig. 1). While historic stock numbers are not well documented, some estimates suggest approximately 30,000 and 1,500 whales inhabited the east and west, respectively, prior to the height of their exploitation (Scammon, 1874; Rice and Wolman, 1971). Both stocks were severely depleted by commercial whaling, and rapid population declines occurred from 1846-1900 in the east Pacific and 1910-1933 in the west Pacific (Rice and Wolman, 1971). By 1933, western gray whales were considered extirpated by some authorities (Bowen, 1974). After commercial whaling ceased due to international agreements in 1937 and 1946, whales in the east rebounded to approximately 27,000 individuals by the late 1980s (Eguchi, 2023). Additionally, a small group of roughly 200 whales appeared in the western Pacific near Sakhalin Island, Russia (Rice and Wolman, 1971) (Fig. 2).

### Estimated Gray Whale Population Changes (1840-2000)

Several factors complicate the designation of the WNP whales as western gray whales. Gray whales complete seasonal migrations that are some of the longest known of any vertebrate, and at 22,511 km, an individual gray whale holds the record for the longest known round-trip mammalian migration (Mate et al., 2015). The species is known to travel long distances even outside of these migratory routes, and in recent years a gray whale has been found in the Atlantic (Hoelzel et al., 2021) and in the Mediterranean (Scheinin et al., 2011). Thus, the intrusion of ENP gray whales into the western Pacific is a reasonable possibility. This is especially true if a) gray whales routinely migrated between the eastern and western Pacific and individuals slowly “returned” to the western Pacific upon the cessation of commercial whaling or if b) individuals in the east, where population size rapidly expanded in the post-commercial whaling era, moved west in search of untapped food sources or less competition. In addition, the range of the historical western stock has never been well characterized, and the exact location of their wintering grounds is unknown, though it was postulated to be somewhere in the South China Sea (Andrews, 1914; Weller and Brownell, 2012). Contemporary WNP whales are not known to winter south of Sakhalin Island, and in recent decades only a few gray whale sightings have been recorded in these waters (Cooke et al., 2017; Nakamura et al., 2019). By contrast, WNP whales have been documented migrating to the eastern Pacific; from 2010-2011, a tracking study followed three whales tagged in the WNP near Sakhalin Island. All three migrated east before the tracker fell off two whales, but the third was tracked to the eastern gray whale wintering grounds in Baja California, Mexico, and then back to Sakhalin Island, Russia (Mate et al., 2015).

Genetic and genomic analyses have provided mixed evidence regarding differentiation between ENP and WNP whales. Subsequent genomic studies based on panels of single nucleotide polymorphisms (SNPs) revealed only modest differentiation between whales sampled in the ENP and those sampled in the WNP, suggesting mixed-stock aggregations in both the Western and Eastern Pacific (Brüniche-Olsen et al., 2018a; DeWoody et al., 2017). However, recent microsatellite work indicates that while gene flow occurs between gray whales sampled in the ENP and the WNP, private alleles in the WNP are maintained by matrilineal site fidelity and non-random mating (Lang et al., 2022). Despite this evidence of differentiation at biparentally-inherited nuclear markers, mitochondrial haplotypes are shared between whales sampled in the WNP and the ENP (Brüniche-Olsen et al., 2021). Collectively, these genetic and genomic studies indicate a lack of reproductive isolation that is manifested as weak contemporary structure between the ENP and WNP. From a conservation perspective, the more important question is whether this observed structure is due to a recolonization (as argued by Bickham et al., 2020) or the cryptic persistence of a remnant western gray whale lineage (Cooke et al., 2020).

Many scenarios have been generated, but not tested, to explain the evolutionary origins of WNP whales and the modern-day population structure of the Pacific gray whale (Punt et al., 2025). Our goal was to critically evaluate the genomic ancestry of gray whales sampled in the WNP using the high-resolution data offered by whole genome resequencing. We present a population genomics study of gray whales sampled from both the eastern and western Pacific, with whole genome resequencing data for 74 individuals. Our analyses offer new insights into both historical divergence and present-day structure, providing critical context for conservation and management.

## Methods

### Sampling, DNA extraction, and whole genome resequencing

We collected tissue biopsy samples between 2011 and 2016 via a compound crossbow with a 150 lb draw weight and 40mm by 7mm internal diameter tip arrows. The samples in this study were previously used for SNP panel and mitochondrial genome analyses, and three whole-genome sequences were generated in previous studies (Brüniche-Olsen et al., 2018a,b; DeWoody et al., 2017; Brüniche-Olsen et al., 2021). We performed DNA extractions on tissue biopsies from an additional 30 ENP and 39 WNP gray whales. Whole-genome sequencing was done using an Illumina NovaSeq to produce 150bp paired end reads. Additionally, we included short read sequencing from two published eastern gray whale whole-genome SRAs (SRR5665641 and SRR5665642; Árnason et al., 2018). We confirmed short read quality with FASTQC (v. 0.12.1; Andrews, 2010) and mapped to a chromosome-level gray whale genome assembly (GCA 028021215.1) using the bwa mem algorithm (v. 0.7.17; Li, 2013). Ambiguous regions were identified with GenMap (v. 1.3.0; Pockrandt et al., 2020). Repeats were identified with RepeatMasker using the “mammals” repeat library (v. 4.1.2; Smit et al., 2015). After mapping, we used GATK 3.8 to remap around indels (McKenna et al., 2010). Unmapped, secondary, QC failed, duplicate, supplementary, and unproperly paired alignments were removed with SAMtools (v. 1.17; Li et al., 2009). The BAM files obtained from these steps were used for subsequent analysis.

### Population structure and phylogenomics

For analyses of population structure, we calculated genotype likelihoods and allele frequencies using ANGSD (v. 0.940; Korneliussen et al., 2014) with minimum base and mapping quality filters (minQ = 20; minMapQ = 20), SNP calling with a likelihood ratio test (SNP_pval = 1×10⁻⁶) and a minor allele frequency threshold of 0.05. Analysis-specific parameters (e.g., depth filters, major/minor allele inference, IBS/HWE calculations) differed depending on the downstream method, and full scripts can be found on GitHub (https://github.com/nataliemallen/Gray_whales/tree/main).

To help identify close relatives such as cow/calf pairs, we used allele frequencies to calculate pairwise relatedness with NgsRelate (v. 2; Korneliussen and Moltke, 2015). Pairwise relationships were inferred using a combination of R1-KING statistics (Waples et al., 2019) to identify first degree (full siblings/parent-offspring), second degree (half-siblings/avuncular/grandparent-grandchild), third degree (first cousins), and unrelated pairs (Fig. S1). First-degree relationships were distinguished from full siblings versus parent-offspring using the R1 statistic. To visualize population structure without a null model, we used genotype likelihoods to calculate a covariate matrix using PCAngsd (Meisner and Albrechtsen, 2018) as part of a Principal Component Analysis (PCA). To examine structure more closely, we also used the same method to calculate covariate matrices independently for each sample site and for each chromosome. To investigate the possibility of unique genomic structural variants in the ENP and WNP, we ran PCAngsd in non-overlapping windows of 100,000 mb across the genome and plotted sliding window PCAs for each chromosome (Ravagni et al., 2025). Admixture plots for K from 1 to 5 were also generated based on a model of Hardy-Weinberg equilibrium using genotype likelihoods in NgsAdmix (Skotte et al., 2013). To examine the most likely number of clusters (i.e., genetic populations), the Evanno method was used to calculate ΔK (Evanno et al., 2005). Because this method cannot detect K = 1, we also calculated correlations of the residuals using evalAdmix (Garca-Erill and Albrechtsen, 2020) to evaluate model fit. For comparative metrics heavily impacted by sample size, we randomly downsampled WNP individuals to n=33 to match our ENP sample size; we hereafter refer to this sampling of n=66 total individuals as the downsampled dataset. We used the downsampled dataset to calculate pairwise *F*_ST_ and *D*_XY_ between ENP and WNP with ANGSD. Standard error for global pairwise *F*_ST_ was estimated using a block jackknife over 500kb windows. Given our extremely large number of windows, we evaluated the statistical significance of *F*_ST_ estimates by sampling subsets of 100-50,000 non-overlapping windows with 1,000 replicates each.

A nuclear phylogeny was generated from our genome sequences to infer evolutionary relationships between populations. We mapped short reads from a fin whale (*Balaenoptera physalus*; SRR23615109) to use as an outgroup and called genotypes with a minimum depth of 10X using ANGSD (Korneliussen et al., 2014). The resulting VCF file was used to obtain maximum likelihood trees with 1000 bootstraps in IQTREE2 (v. 2.2.2.9; Minh et al., 2020). In addition to this tree, we estimated genetic relationships among individuals with an identity-by-state (IBS) approach that does not rely on genotype calls. We generated pairwise IBS estimates using ANGSD (Korneliussen et al., 2014) and inferred a neighbor-joining tree with FastME (Lefort et al., 2015). To further assess population structure using the IBS distance matrix, we conducted a principal component analysis (PCA) and used hierarchical clustering to compare pairwise IBS values among and within sample sites.

We examined copy number variants (CNVs; Jakobsson et al., 2008) because they may reflect population structure more effectively than SNPs (Dorant et al., 2020). To compare CNVs between ENP and WNP samples, we used the downsampled dataset. Copy number variants were identified using CNVpytor (v1.3; Abyzov & Urban, 2020) with a custom configuration optimized for *E. robustus*. For each sample, read-depth histograms were generated at 10 kb and 100 kb bin sizes, followed by GC correction, regional partitioning, and CNV calling. Population-level CNV frequencies were compared between ENP and WNP using Fisher’s exact test implemented in Python v. 3.11.5. Contingency tables were constructed from counts of individuals with and without each CNV in the two populations to identify significant frequency differences. Overall CNV abundance per individual was compared between populations using a Wilcoxon Rank-Sum Test. Differences in the proportion of duplications versus deletions were assessed with a chi-square test of independence.

### Allele frequency spectra and neutrality tests

For each sample site, we estimated site frequency spectrum (SFS) and Tajima’s D using ANGSD and the downsampled dataset. We inferred the SFS from site allele frequency likelihoods using realSFS with folded spectra to account for unknown ancestral states. The SFS was then converted to thetas with saf2thet, and Tajima’s D values (genome-wide and sliding-window) were computed using thetaStat with a window size of 50 kb and a step size of 10 kb. To evaluate differences in the SFS between sample sites, we used two statistical approaches. A two-sample Kolmogorov-Smirnov (KS) test was used to compare the overall SFS distributions. Differences in Tajima’s D between populations were assessed using a two-sided Wilcoxon rank-sum test in R. Effect size was evaluated as the difference in medians, and uncertainty in the mean estimates was quantified by bootstrapping (10,000 replicates) to generate 95% confidence intervals.

### Genomic diversity

We calculated genomic diversity from genotype likelihoods using individual genome-wide heterozygosity, which we defined as the number of heterozygous sites divided by the total number of sites analyzed in the genome and calculated using genotype likelihoods. We compared mean heterozygosity (*H*) to existing data in related marine mammals including other whale species impacted by commercial whaling. A Wilcoxon sum-rank test was used to test for a significant difference between mean heterozygosity in the ENP versus WNP. Associations between heterozygosity and sequencing breadth or depth were assessed using linear regression.

### Analysis of individuals based on admixture coefficients

To test whether admixture is recent (<5 generations), we assessed the relationship between genome-wide heterozygosity and admixture coefficients in WNP individuals using Spearman rank correlation. Recent admixture between differentiated populations would be expected to elevate heterozygosity in admixed individuals. We calculated expected admixture coefficient ranges for F1 hybrids (0.50 ± 3 standard errors) and first-generation backcrosses (0.25 and 0.75 ± 3 SE) using a simple binomial sampling model across the 3,026,989 sites used to estimate admixture. This approximation assumes independent sites and fixed allele frequencies and thus uncertainty in genotype likelihoods and deviations from assumed historical allele frequencies may inflate variance around these expectations. Nonetheless, given the large number of sites, the uncertainty of admixture proportions is extremely small and our power to detect recent hybrids is high. Finally, we calculated genome-wide *F*_ST_ using the methods above between 7 ENP individuals and 7 WNP that appeared to have very little eastern ancestry (individuals furthest to the right on the PCA; Fig. 3) to quantify the most extreme differences between ENP and WNP whales.

**Figure 3:**
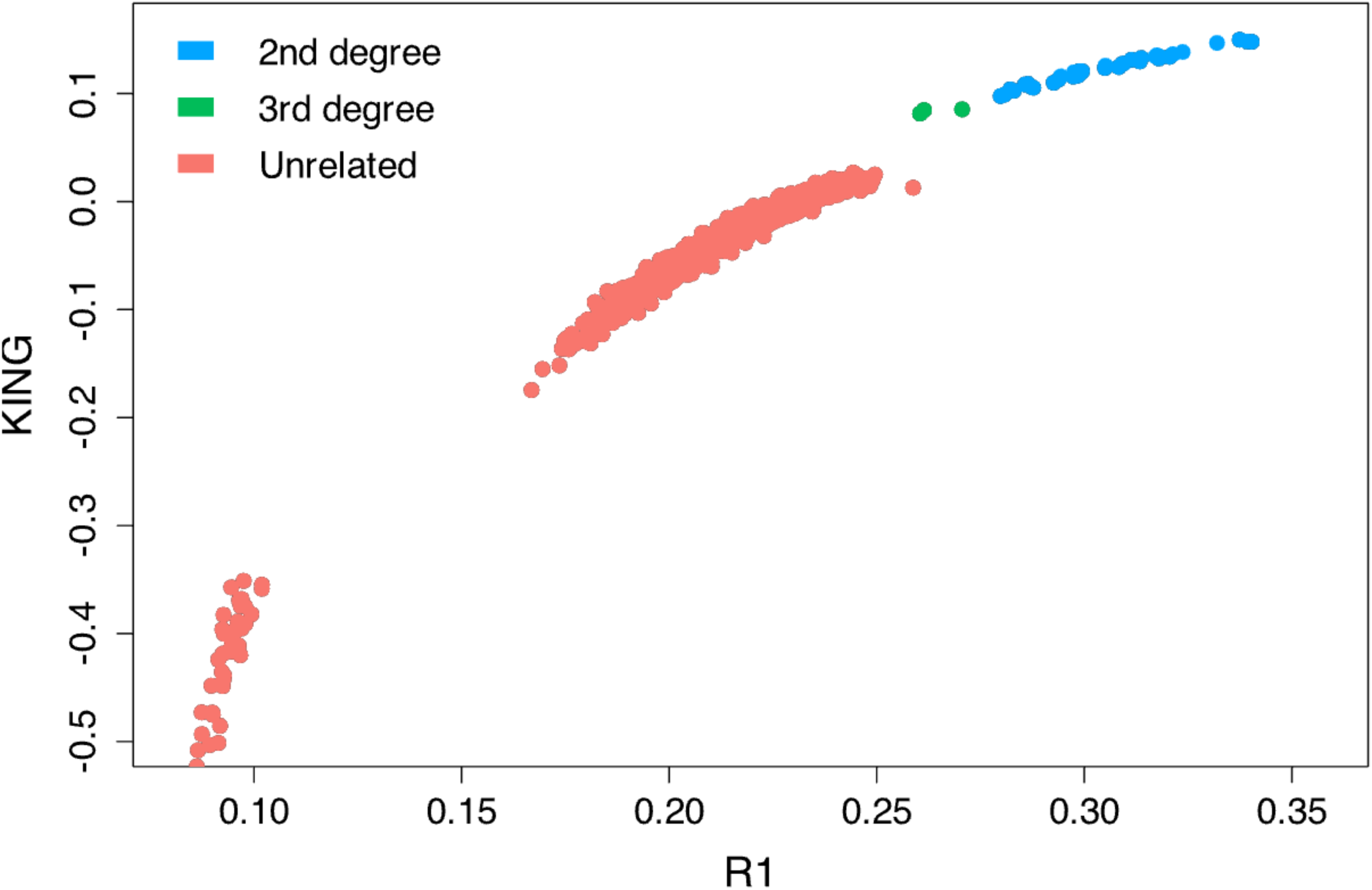
*Kinship plot including* only *pairs of individuals from different sample sites. No pairs of first-degree relatives were identified at different sites, but several second- and third-degree relatives were.* Alt text: Scatterplot with R1 on the x axis and KING on the y axis. Second degree relative pairs are shown as blue dotes, while third degree relative pairs are shown as green dots and unrelated pairs are shown as red dots. The dots follow a curved path along the plot and the majority of dots are red. There are three green dots and approximately 20 blue dots.

## Results

### Sequence data and quality control

Across our 74 samples, average breadth and depth of coverage across the genome after mapping and quality control were 61.3% and 5.48X, respectively. Breadth of coverage was in the expected range as 32.8% of the reference genome was repeat-masked. Depth of coverage was variable and ranged from 1.7X to 29.3X, but most samples fell within the range of 3-5X. Average depth was 5.3X in ENP samples and 5.6X in WNP samples. SNP counts used for each analysis differed by filtering parameters, but most genotype likelihood approaches used over 3 million sites (for example the PCA included 3,311,553 SNP sites).

### Population structure and phylogenomics

During sample collection in the field, 11 potential mother-calf pairs were identified by field biologists, and our kinship analysis corroborated 10 of these. All pairs of putative first-degree relatives were sampled from the same geographic locations and thus plausible. While most pairs sampled from different locations were unrelated, we identified 38 second-degree and 3 third-degree pairwise relationships in individuals sampled from different locations (Fig. 3).

Principal component analysis of all individuals revealed a pattern of ENP individuals clustering within WNP individuals, with a striking degree of variation in the west (Fig. 4B). We carefully examined the possibility of technical factors influencing this result but found none that changed the pattern, including when regions of Linkage Disequilibrium (LD) were removed and when all first, second, and third-degree relatives were excluded (Fig. S2). This structure was consistent in PCA plots for each sample site and each chromosome (Figs. S3 & S4). Sliding window PCAs for each chromosome did not reveal any structural variants that differentiated gray whales collected at the different sample sites (Fig. S5). Admixture analysis best supported K = 2 (Table S1), and structure plots show much more variation in the WNP than the ENP (Fig. 4A; Fig. S6). Pairwise correlations of residuals generated with evalAdmix were closer to zero at K = 2 than K = 1, indicating a better model fit at the higher K value (Fig. S7). This is contrary to what would be expected in the case of panmixia. The phylogenomic tree displays notable structure, with all but four individuals sampled from the WNP clustering together (Fig. 4A).

**Figure 4:**
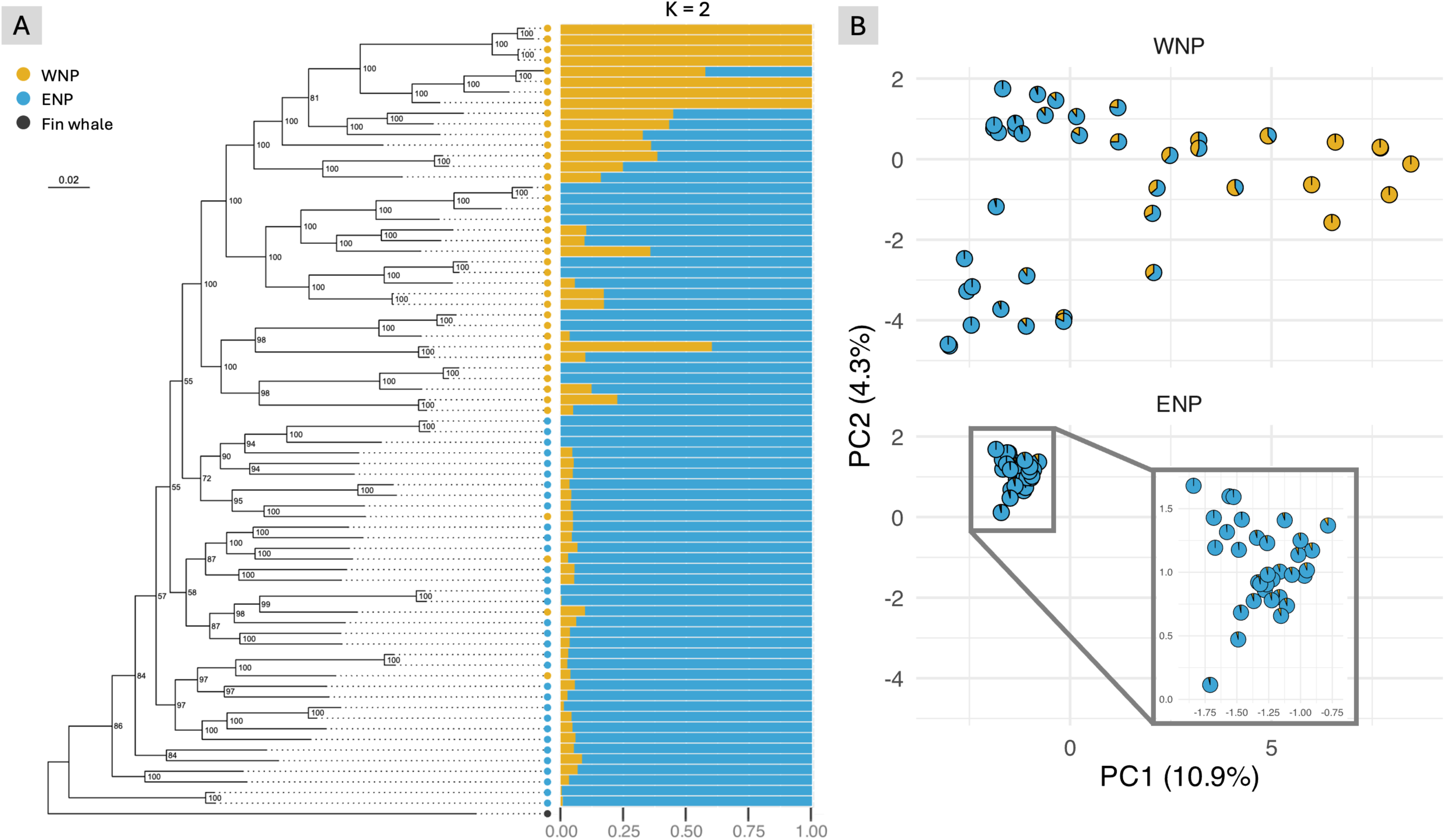
*Population structure in the Pacific gray whale. Phylogenomic tree, rooted with fin whale (*Balaenoptera physalus*), where samples collected from the ENP are in blue and samples collected from the WNP are in yellow. Admixture plots for K=2 are organized in order of tree branch tips. On the right, the PCA is faceted by sampling site and each individual is plotted with K=2 admixture coefficients as a pie. Across population structure analyses, there is more variation in individuals from the WNP than individuals from the ENP.* Alt text: Panel A shows a phylogenomic tree with ENP individuals in blue and WNP individuals in yellow. Most yellow dots are at the top of the plot, nested within the clade of blue dots, but four yellow dots are scattered among the blue. Aligned with each branch tip is a K=2 admixture plot in blue and yellow. The top four bars in the plot are solid yellow, followed by one bar about half yellow, followed by 3 solid yellow bars. The next seven bars are between 10-50% yellow. The rest of the bars, moving down the tree, are mostly blue, with a few scattered bars that are 10-50% yellow matching yellow individuals on the tree. The bottom half of the bars, which match blue individuals, are each about 90-95% blue. Panel B shows the same individuals on a PCA where the top is WNP and bottom is ENP. Each individual is represented as a pie chart of K=2 admixture coefficients. The WNP individuals are much more spread out across the PCA and form a gradient of mostly blue to mostly yellow from left to right. The ENP individuals cluster tightly together on the left side and are all nearly 100% blue. PC1 explains 10.9% of the variation while PC2 explains 4.3%.

We compared results from the genotype and genotype likelihood analyses to those from an IBS distance matrix. Patterns found in the PCA and phylogenomic tree were recapitulated by the IBS approach (Fig. S8). In congruence with the PCA, a plot of IBS distance distributions reveals more variation within whales from the WNP than within whales from the ENP (Fig. 5D).

**Figure 5:**
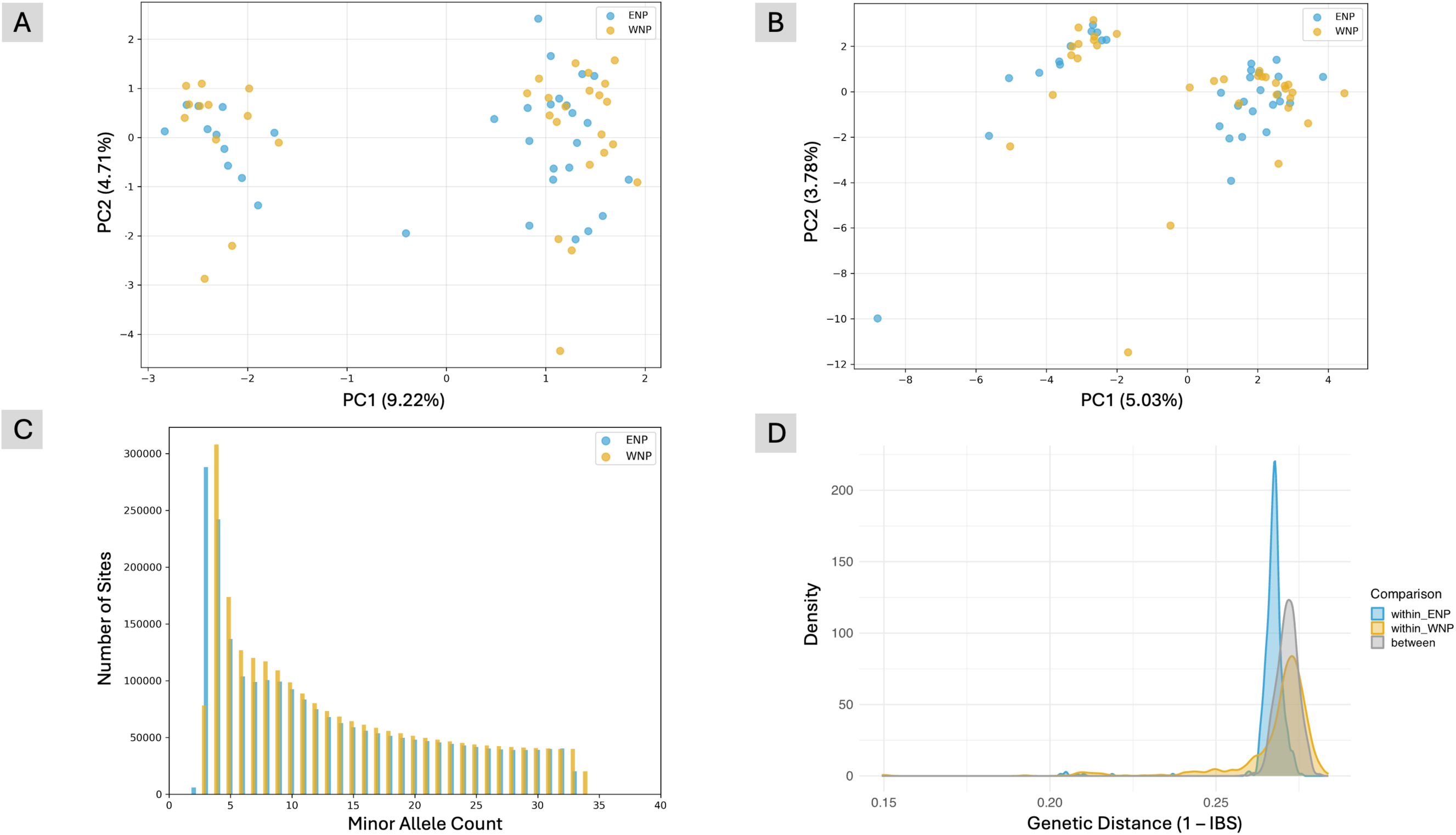
*Comparison of copy number variants, folded site frequency spectra, and Tajima’s D in equal sample sizes of ENP (blue) and WNP (yellow) gray whales. A) PCA based on 1,062 unique CNVs (100 kb resolution). B) PCA based on 5,164 unique CNVs (10 kb resolution). C) Folded Site Frequency Spectrum (SFS) where minor allele count (x-axis) is plotted against the number of sites (y-axis). Bars are offset to avoid overlap. D) Distribution of pairwise genetic distances, calculated using IBS, among and within ENP and WNP. Samples from the ENP and WNP are extremely similar in plots A-C. In plot D, within-population distances are generally lower than between-population distances, but distances within the WNP are higher than those within the ENP.* Alt text: Panel A shows a PCA with two loose clusters, each made up of some ENP and some WNP individuals. PC1 explains 9.22% of variation while PC2 explains 4.71%. Panel B also shows a PCA with two clusters, each made up of some ENP and some WNP individuals. PC1 explains 5.03% of variation while PC2 explains 3.78%. Panel C shows minor allele count on the x axis and number of sites on the y axis. At MAC =4, the number of sites is about 300,000 for both ENP and WNP, and this peak falls sharply to about 70000 by MAC = 15 where it remains relatively steady until MAC=33. The WNP and ENP have very similar distributions, but the WNP is slightly shifted left. Panel C shows a plot of genetic distance (1-IBS) on the x axis and density on the y axis. Within ENP density is a tall, skinny peak, while within WNP density is a shorter wider peak. The “between” density curve is shown in gray and falls between the two within curves.

Weighted genome-wide differentiation between ENP and WNP gray whales was small (*F*_ST =_ 0.019, SE = 2.1e-5, p < 0.001), indicating low but significant differentiation between sampling sites that span much of the North Pacific Ocean. Subsampling analysis of *F*_ST_ gave consistent results regardless of window number, indicating this estimate is robust to sampling variance (Table S2), and global absolute sequence divergence was low (*D*_XY_ = 0.00034).

Principal component analysis of CNVs showed two clusters with mixed sample origin at both 10kb and 100kb resolution (Fig. 5A & B), consistent with a previous SNP panel that indicated mixed-stock aggregations (Brüniche-Olsen et al., 2018a). After correcting for multiple testing, no individual CNVs at either 10kb or 100kb resolution were found to be significantly differentiated between geographic sampling sites based on Fisher’s exact test at α = 0.05. Additionally, no significant differences were detected in overall CNV abundance or type between sampling sites at either resolution (Fig. S9). A marginal difference in the proportion of duplications versus deletions was observed at 10 kb resolution (χ² = 4.90, p = 0.027), though this pattern was not evident at 100 kb resolution (χ² = 2.76, p = 0.097).

### Allele frequency spectra and neutrality tests

We examined SFS and Tajima’s D independently in gray whales sampled from both the WNP and the ENP. Visually, SFS appeared nearly identical between the two, with a very slight right shift in the WNP (Fig. 5C). The KS test indicated no significant difference in the cumulative distributions (D = 0.0896, p = 0.82). These results point to a consistent overall distribution between whales sampled in the ENP and WNP. Mean Tajima’s D was statistically different between ENP and WNP (D_ENP_ = 0.803, D_WNP_ = 0.816; Wilcoxon rank-sum p < 2.2×10⁻¹⁶; Fig. S10), though the effect size was extremely small (median difference = 0.008). Bootstrapped 95% confidence intervals for the means showed only slight separation between populations (ENP: 2.287–2.291; WNP: 2.278–2.282).

### Genomic diversity

Mean genome-wide heterozygosity, based on 74 genomes was low (*H* = 0.00047; Fig. 6A). Individual heterozygosity showed no significant association with either breadth or depth of sequencing coverage (breadth: p = 0.93; depth: p = 0.12). There was no significant difference in mean heterozygosity between whales sampled from the WNP and the ENP (W = 593, p = 0.3689). Our data indicate gray whales have relatively low genomic diversity compared to other rorquals (Árnason et al., 2018; Fig. 6B).

**Figure 6:**
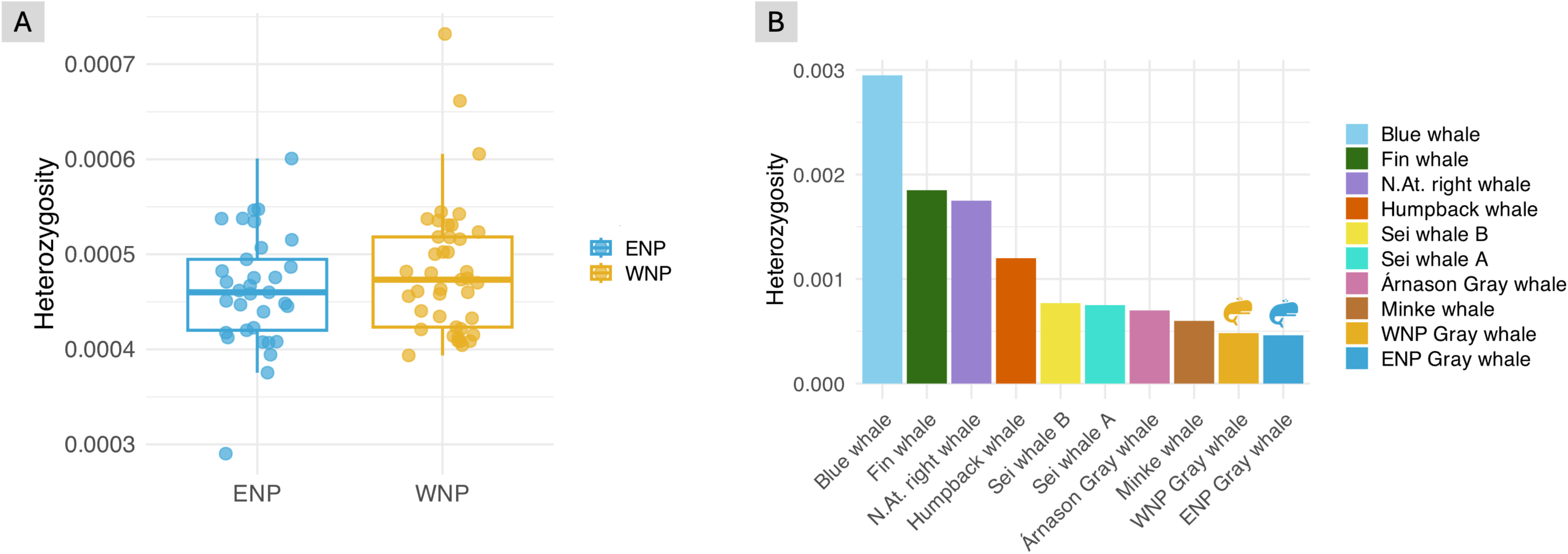
*Genome-wide heterozygosity. A) Box plots of individual genome-wide heterozygosity with ENP individuals in blue and WNP in yellow. B) Comparison of our gray whale heterozygosity estimates in blue and yellow (far right; denoted by whale icons) compared to estimates by Árnason et al., (2018) of gray whale and other rorquals. Samples from the ENP and WNP have very similar heterozygosity, and gray whales generally have lower heterozygosity compared to other rorquals (Blue whale, Fin whale, Humpback whale, Minke whale, and Sei whale).* Alt text: Panel A shows box plots in blue for the ENP and yellow for the WNP with Heterozygosity on the y axis. Raw data points are visible over the box plots. The two box plots have similar means around 0.0047, though the ENP has a lower minimum and the WNP has a higher maximum. Panel B shows a bar chart with baleen whale species on the x axis and heterozygosity on the Y axis. From left to right, blue whale has the highest value (0.00295), followed by fin whale (0.00185), North Atlantic right whale (0.00175), and humpback whale (0.0012). Intermediate values include Sei whale A (0.00075) and Sei whale B (0.00077), minke whale (0.0006), and the gray whale estimate reported by Árnason (0.0007). The ENP gray whale (0.000462) and WNP gray whale (0.000483) have the lowest heterozygosity values in the plot and these two have a whale icon over the bar to indicate the data is from this study.

### Analysis of individuals based on admixture coefficients

We did not identify any significant relationship between genome-wide heterozygosity and admixture coefficients in WNP individuals (Spearman’s ρ = −0.166, p = 0.30; Fig. S10). We also did not find any WNP individuals with admixture coefficients consistent with F1 hybrids or first-generation backcrosses (Fig. S11). Weighted genome-wide *F*_ST_ between 7 ENP individuals and 7 WNP that appeared to have very little eastern ancestry was 0.115, which is much higher than the F_ST_ of 0.019 we found between WNP and ENP overall.

## Discussion

The ambiguous gene flow dynamics of highly mobile marine species are a challenge to evolutionary and conservation biologists. In species such as the gray whale, where commercial whaling imposed recent extreme bottlenecks, these patterns are further obstructed while questions related to conservation are more dire. Researchers have long struggled to discern between historical lineage persistence and recent recolonization of gray whales in the western north Pacific (Scordino et al., 2024). We investigated this question using a comprehensive whole-genome dataset from 74 individuals, providing the most extensive genomic assessment of population connectivity in gray whales to-date. By examining population structure across the Pacific, we aimed to clarify whether whales summering in the west represent remnants of a historical western lineage, recent migrants from the east, or a combination of both. We provide new insights into how migratory marine mammals recover after severe population declines, and our results speak to the power of genomic data to clarify patterns of gene flow in highly mobile marine organisms.

Our genomic analyses reveal contemporary connectivity in the North Pacific rather than isolation between the ENP and WNP. The presence of second- and third-degree relatives sampled at different locations is consistent with tracking studies by Mate et al., (2015), which documented WNP individuals migrating east in the winter. Despite this cohesion, WNP individuals have greater within-group variation than ENP individuals, indicating that admixed WNP individuals harbor some distinct ancestry from ENP individuals (Fig. 4B). The asymmetric pattern of admixture, characterized by minimal WNP ancestry in the ENP, suggests that gene flow has been unidirectional from east to west (Fig. 4A). This scenario is consistent with both the relatively small number of WNP whales and the demographic expansion of ENP whales over the last three decades. The mean overall genomic differentiation between WNP and ENP whales is small relative to a subset of the most differentiated ENP and WNP individuals. The most parsimonious underlying process explaining this pattern is that some western gray whale ancestry persisted through the whaling bottleneck and is now detectable only as ghost introgression within contemporary WNP individuals.

While our results support the persistence of historic western gray whale ancestry in the WNP, they also indicate that much of this signal has been swamped by extensive gene flow from the east. Multiple metrics of diversity and demography were very similar the ENP and WNP, suggesting admixture has homogenized variation across the north Pacific. The difference in Tajima’s D, for example, was extremely small, and while this was statistically significant due to the large number of markers employed, the biological significance (if any) is likely negligible. Similarly, SFS distributions were not statistically different, though the slight right shift visible in the SFS of WNP whales indicates a small deficit of rare alleles that may be due to a more extreme recent bottleneck or smaller long-term effective population size (*N_e_*) compared to ENP whales. Likewise, genome-wide heterozygosity and CNVs showed no clear regional differentiation, and if whales in the ENP and WNP had entirely different demographic histories, the much larger *N_e_* of the historical ENP stock should allow it to more effectively purge harmful CNVs through selection (Cheeseman et al., 2016). Despite this overall homogenization, we found some evidence pointing to the possibility that the Sakhalin Island summering grounds historically harbored a mixed-stock aggregation of whales, with some eastern gray whales making the long migration to Russia each year. Principal component analyses of CNVs have two mixed-stock clusters, consistent with previous SNP data on some of the same individuals (Brüniche-Olsen, et al., 2018a). Variation in the distance of ENP whale migrations is well-documented; the Pacific Coast Feeding Group summers along the coasts of northern California, Oregon, and Washington rather than continuing north to the Bering Sea (Lagerquist et al., 2019). Overall, these results suggest that extensive admixture from the east has largely homogenized genomic variation across the Pacific such that ghost ancestry from the historical western lineage is only faintly detectable, even with high-resolution data.

We further argue that introgression between the ENP and WNP is unlikely to be a solely contemporary phenomenon driven by the aftermath of commercial whaling. Rather, we think it is probable that some degree of historical admixture existed across the Pacific prior to commercial whaling. After the western gray whale stock was feared extirpated in 1933, three gray whales thought to be members of the ENP stock were captured in the Western Pacific in 1942, 1959, and 1968 (Bowen, 1974). Assuming reproductive isolation prior to whaling and an approximately 18-year generation time (Rice and Wolman, 1971; Heppell et al., 2000), at most five generations have passed since ENP whales could have begun traveling to Sakhalin Island. We would expect elevated heterozygosity in recently admixed WNP individuals due to an influx of new alleles (Reeves et al., 2025), but we found no correlation between heterozygosity and admixture coefficients. We also found no individuals with admixture coefficients in the expected ranges for F1 hybrids or F2 backcrosses. Given that admixture analyses were conducted with over 3 million SNPs, we have extremely high power to detect hybrids, and their absence points to some level of background gene flow occurring across the Pacific Ocean prior to 1933. The relatively high *F*_ST_ between ENP and non-admixed WNP individuals indicates such gene flow may have been minimal, though the possibility of a historical mixed-stock aggregation in the west Pacific cannot be discounted and could be responsible for seemingly conflicting results across microsatellite, mitochondrial, and genomic datasets and from previous studies (Lang et al., 2022; Brüniche-Olsen et al., 2021; Brüniche-Olsen et al., 2018b). In the last century, gene flow from ENP to WNP has likely increased dramatically, leading to the patterns of population structure we find in whales sampled from the WNP. The preponderance of evidence presented here and in previous studies (Brüniche-Olsen et al., 2018a; DeWoody et al., 2017; Lang et al., 2022) indicates that some individuals in the WNP harbor unique ancestry compared to the ENP, and the most plausible source of this admixture is historical western gray whale ancestry.

### Limitations and future directions

As with other presumed extirpations, definitive evidence that individuals from the WNP contain western gray whale ancestry would require historical western samples from the pre-whaling era. Such samples are difficult to acquire, and in their absence, we must rely on contemporary genomes. Our data indicates that overall divergence between whales in the ENP and WNP is small due to directional gene flow from east to west, and admixture between the two is likely increasing in the WNP. This is a large step forward in our understanding of gray whale stock structure, though the mostly low coverage sequencing data used here limited our ability to fully reconstruct the evolutionary history of gray whales in the western Pacific. While our results demonstrate that historical gene flow likely occurred across the Pacific, we cannot determine its extent or whether it was directionally balanced. The large influx of recent admixture from east to west makes it difficult to parse historical patterns, particularly in the west. Another possible explanation for our data is historical panmixia, with slow divergence on opposite sides of the Pacific followed by recent homogenization in the west. In a previous study, estimations of historical demography for one sample from the ENP and two from the WNP were similar but did not support ancient panmixia (Brüniche-Olsen, et al., 2018b). All three whales from that study were included in our dataset, but we found both WNP individuals to be admixed, and thus high-coverage sequencing of more individuals is necessary to reconstruct demographic histories and estimate divergence times between examine historical population dynamics in the Pacific. Higher-coverage data would enable reconstruction of demographic histories for more individuals from each sample site, as well as the estimation of divergence times between individuals thought to descend from eastern and western gray whales. Such data could also be used to characterize adaptive genetic variants in WNP whales, providing more context for conservation decisions.

### Conservation Implications

From a conservation perspective, our most important finding is that whales in the WNP have mixed ancestry. This complicates the view of contemporary WNP whales as a distinct population, even if they may have been at one time. Our data suggest that naturally occurring admixture from east to west almost certainly prevented the complete extirpation of historical gray whale assemblages in the WNP, allowing ghost introgression to persist. Given recent gene flow and the known long-distance movements of gray whales, future conservation strategies that should benefit the species as a whole might include maintaining migration corridors and preserving foraging habitat rather than focusing on explicit east-west distinctions.

### Conclusions

Our study demonstrates that whole-genome data can help resolve questions about population dynamics in instances of extreme bottlenecks or presumed extirpations. We successfully used whole-genome data to reconcile previous studies of gray whale population structure. The WNP aggregation likely represents a combination of whales with eastern and western ancestry, with admixture occurring between them. We demonstrate that whole-genome datasets provide enough resolution to detect population structure even when rapid introgression has swamped much of the signal. The ability to examine subtle structure is especially relevant in highly mobile species, including many marine mammals, where complex migration and gene flow patterns can be difficult to parse.

## Funding and Acknowledgements

This research was supported in part by Exxon Neftegas Limited and by Sakhalin Energy Investment Company. JAD was supported in part by the National Institute of Food and Agriculture and by the U.S. State Department’s Fulbright U.S. Scholar program. The content herein is solely the responsibility of the authors and does not represent the official views of the funding parties. We thank members of the DeWoody lab for comments on an earlier version of this paper.

## Data Accessibility

Sequencing reads will be made available under SRA Bioproject PRJNA1424656.

