## Supplemental Information for "The contemporary Pacific gray whale (*Eschrichtius robustus*) gene pool includes ancestry from a potential ghost population: inferences from population genomics"

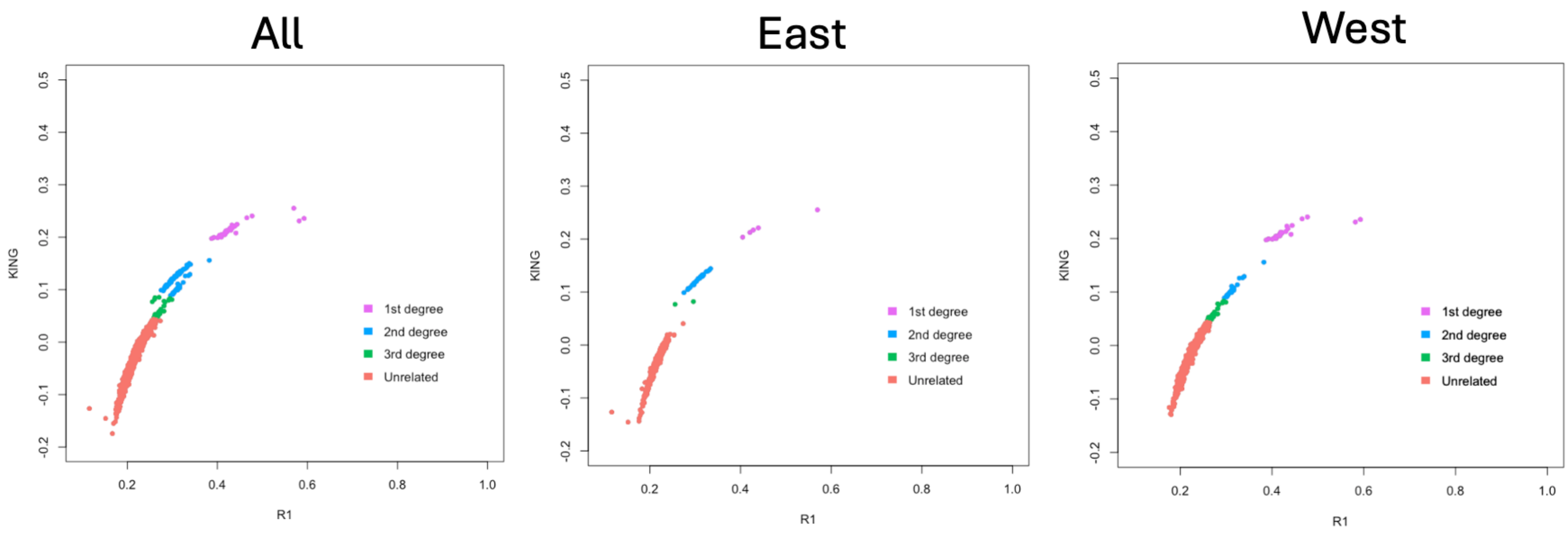


**Supplementary Figure 1:** Kinship plots for our sampled individuals, with all (left), east only (middle), and west only (right).

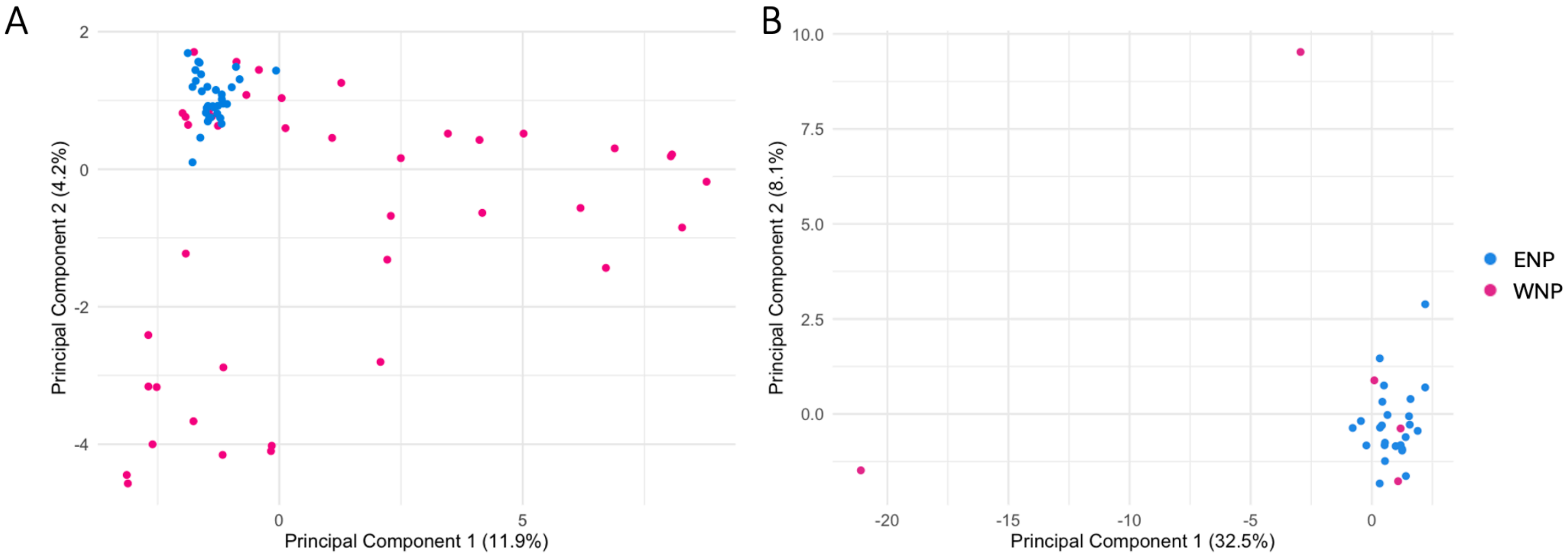


**Supplementary Figure 2:** PCA plots used to examine the possibility of confounding technical factors. A) PCA with LD regions removed. B) PCA with all relatives removed. Removing LD had no effect on the PCA, and removing relatives decreased the number of individuals but still recapitulated the pattern of more spread in the WNP.

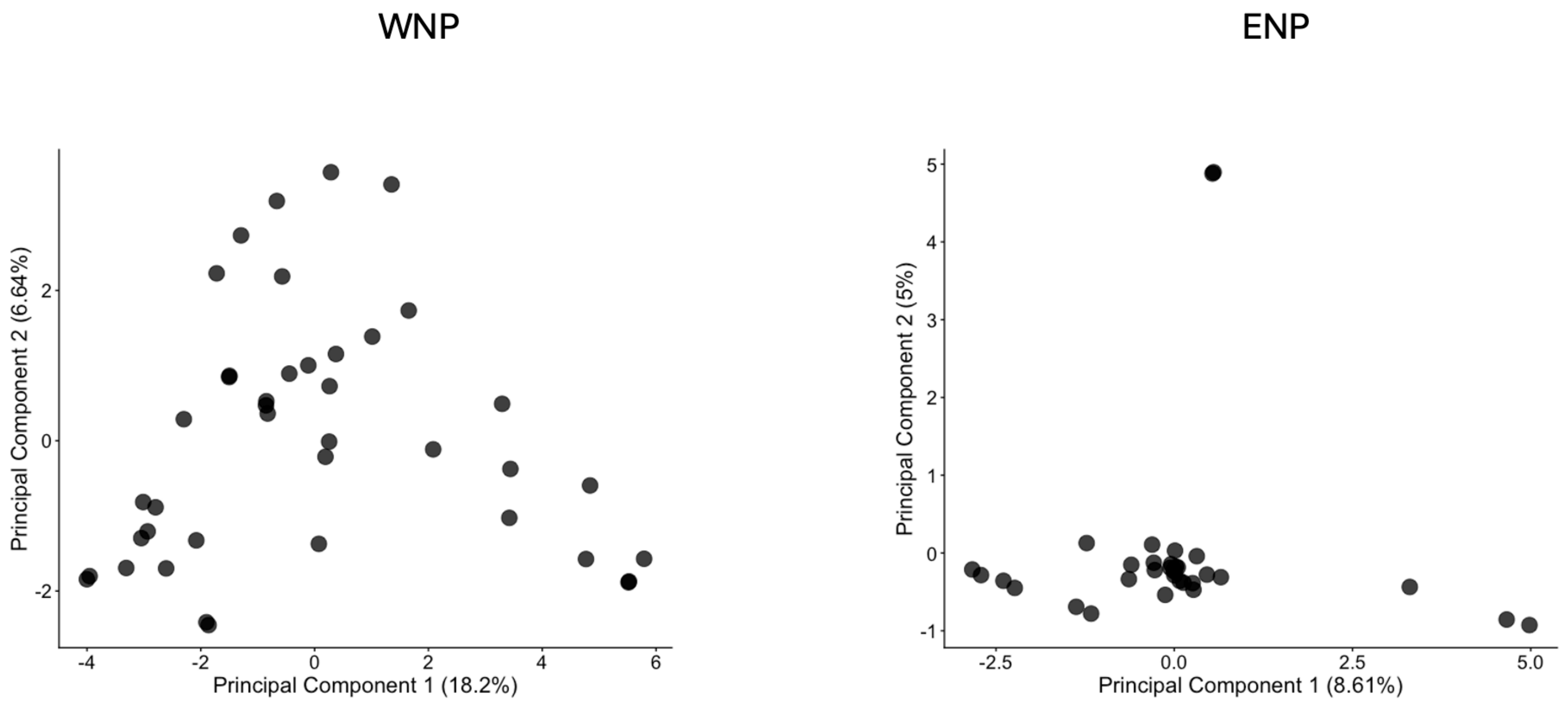


**Supplementary Figure 3:** PCA plots for each sample site.

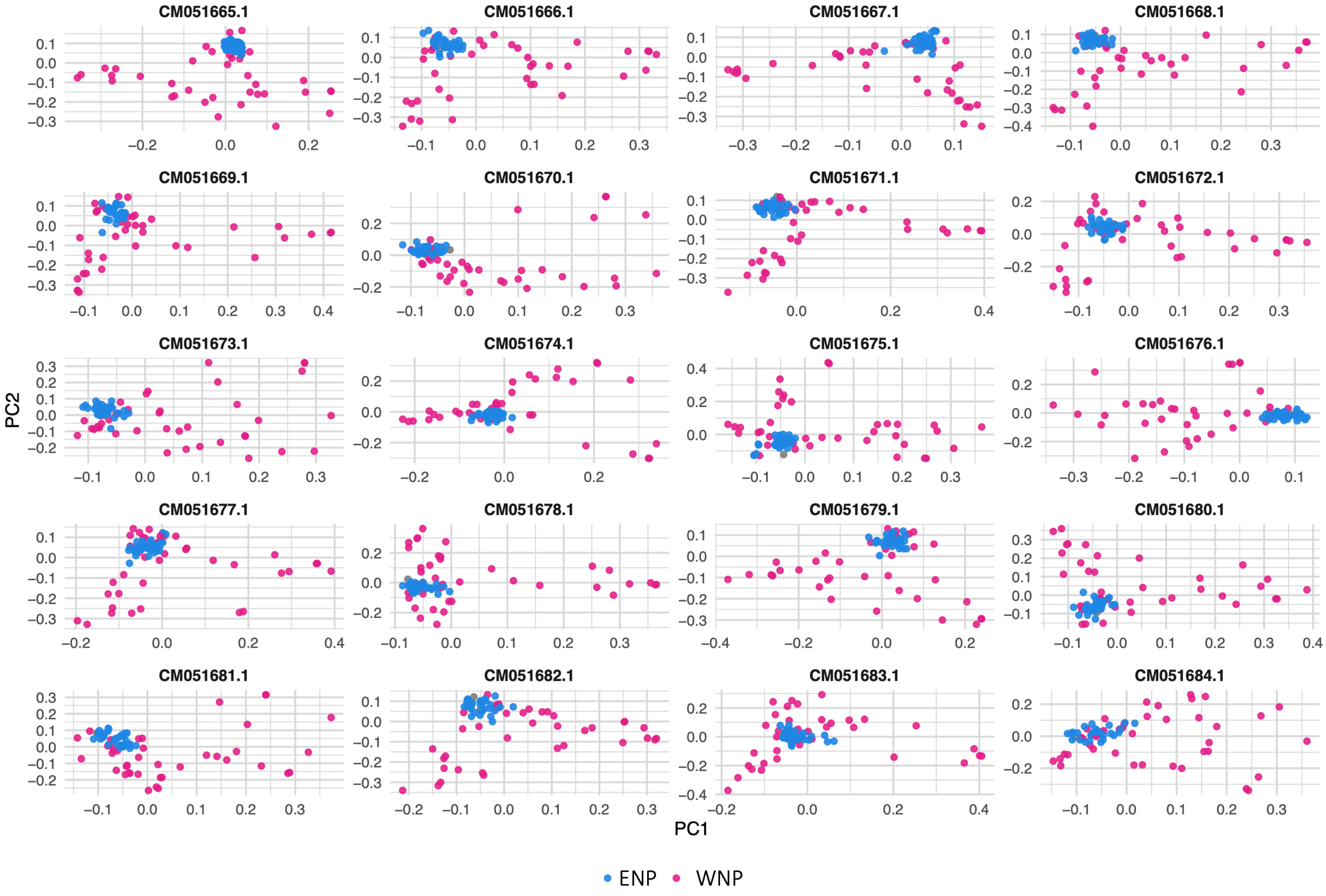


**Supplementary Figure 4:** PCA plots for each chromosome.

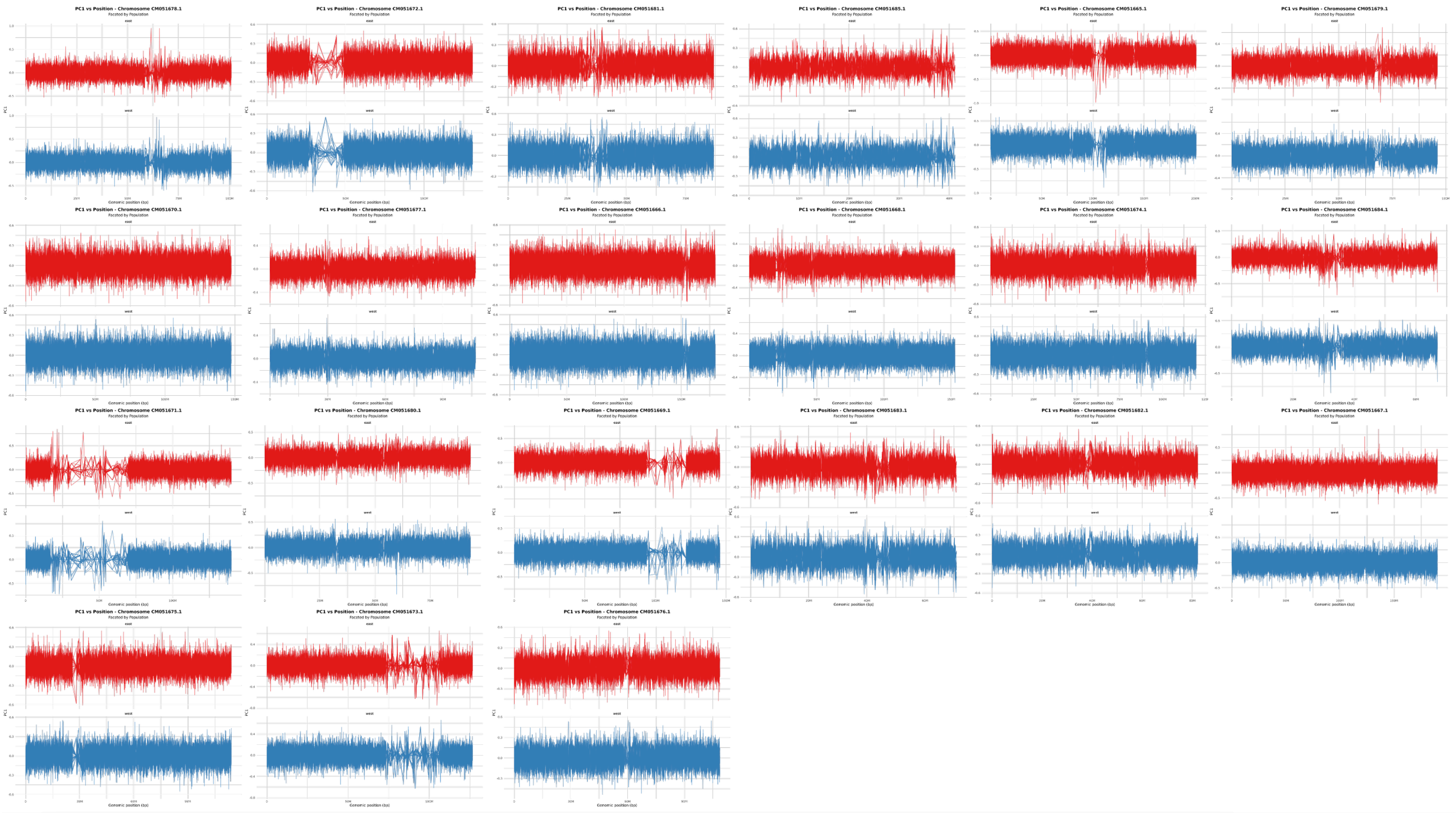

**Supplementary Figure 5:** Sliding window PCA plots for each chromosome. Some regions exhibit patterns characteristic of structural variants, but none of these regions differentiate ENP and WNP whales.

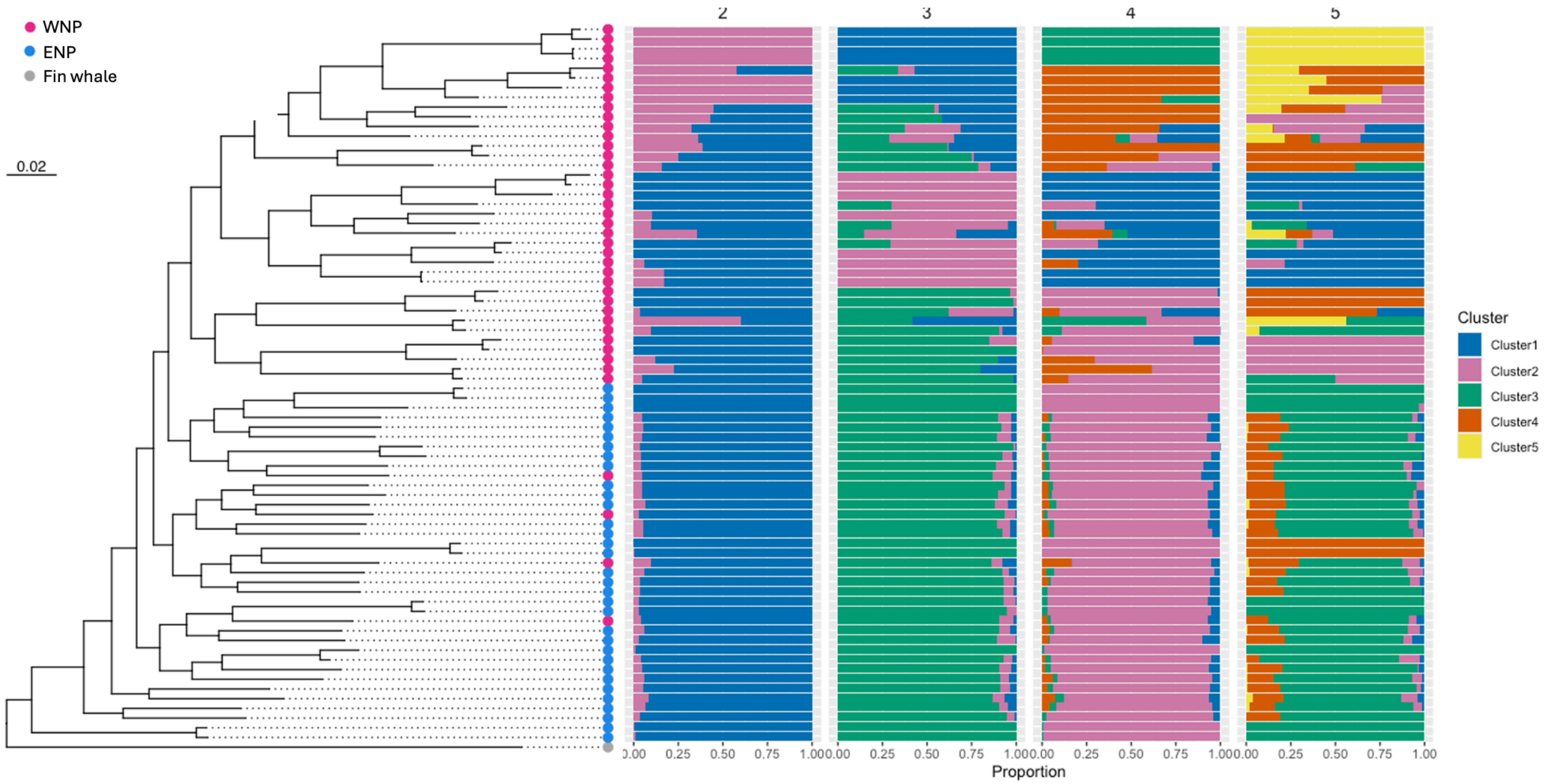


**Supplementary Figure 6:** Phylogenomic tree, rooted with fin whale, where samples collected from the ENP are in blue and samples collected from the WNP are in magenta. Admixture plots for K=2, 3, 4, and 5 are organized in order of tree branch tips.

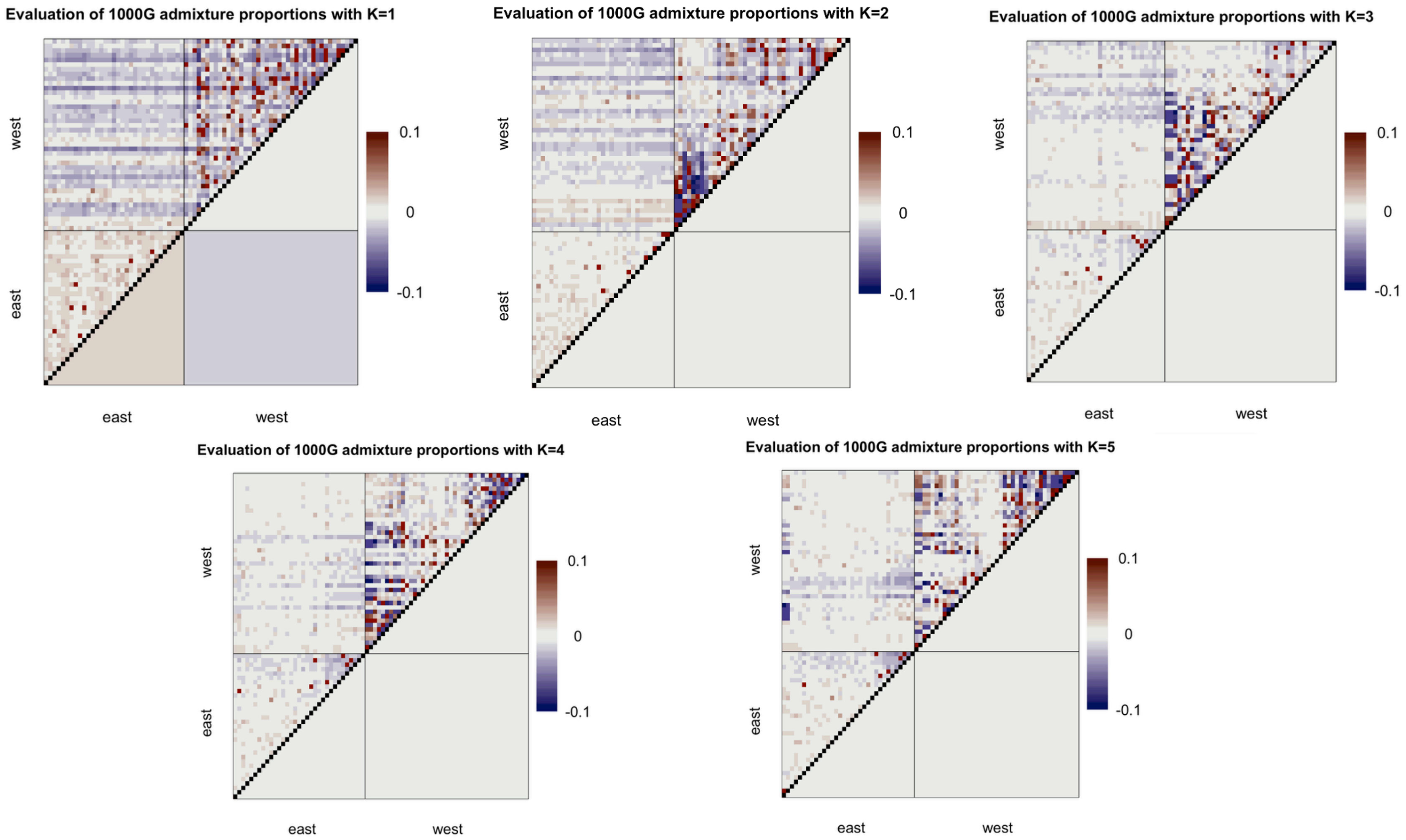


**Supplementary Figure 7:** Plots of evalAdmix pairwise correlations of residuals for K=1-5. More positive correlations are more orange while more negative correlations are more blue; correlations close to zero indicate a better model fit.

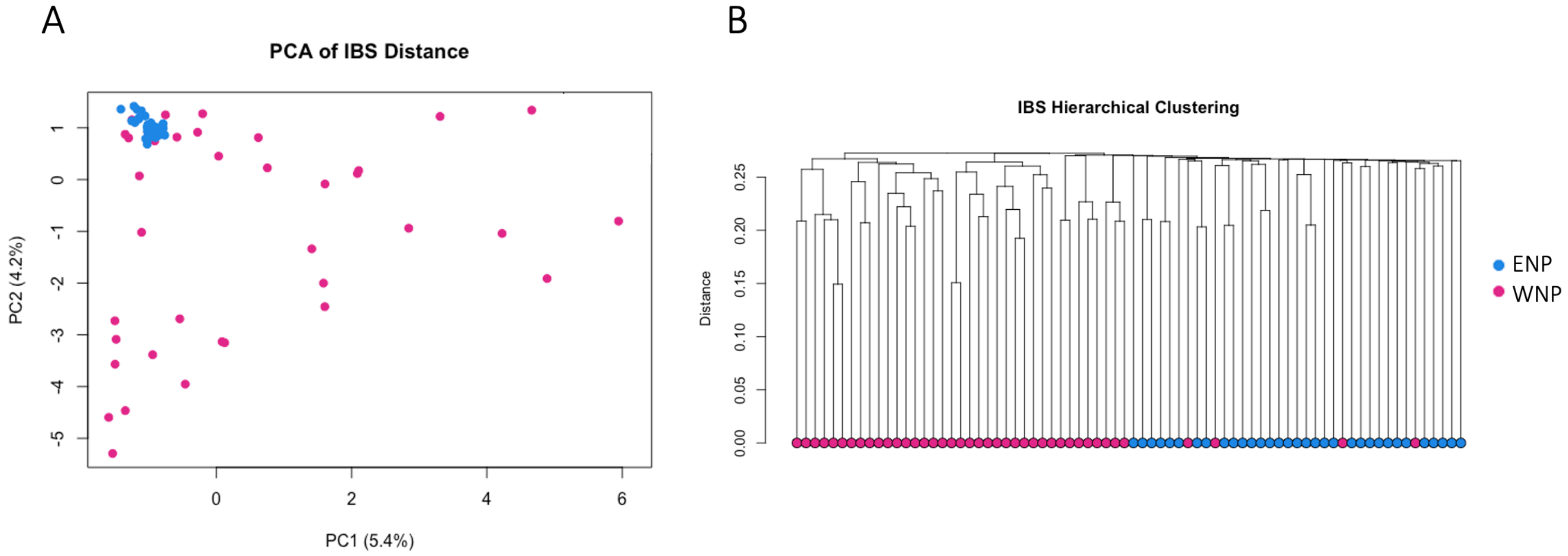


**Supplementary Figure 8:** Identical-by-state (IBS) analysis of population structure. A) PCA of ENP and WNP using IBS distance matrix. B) IBS tree of ENP and WNP individuals. These results recapitulate those found using genotype likelihoods and genotypes.

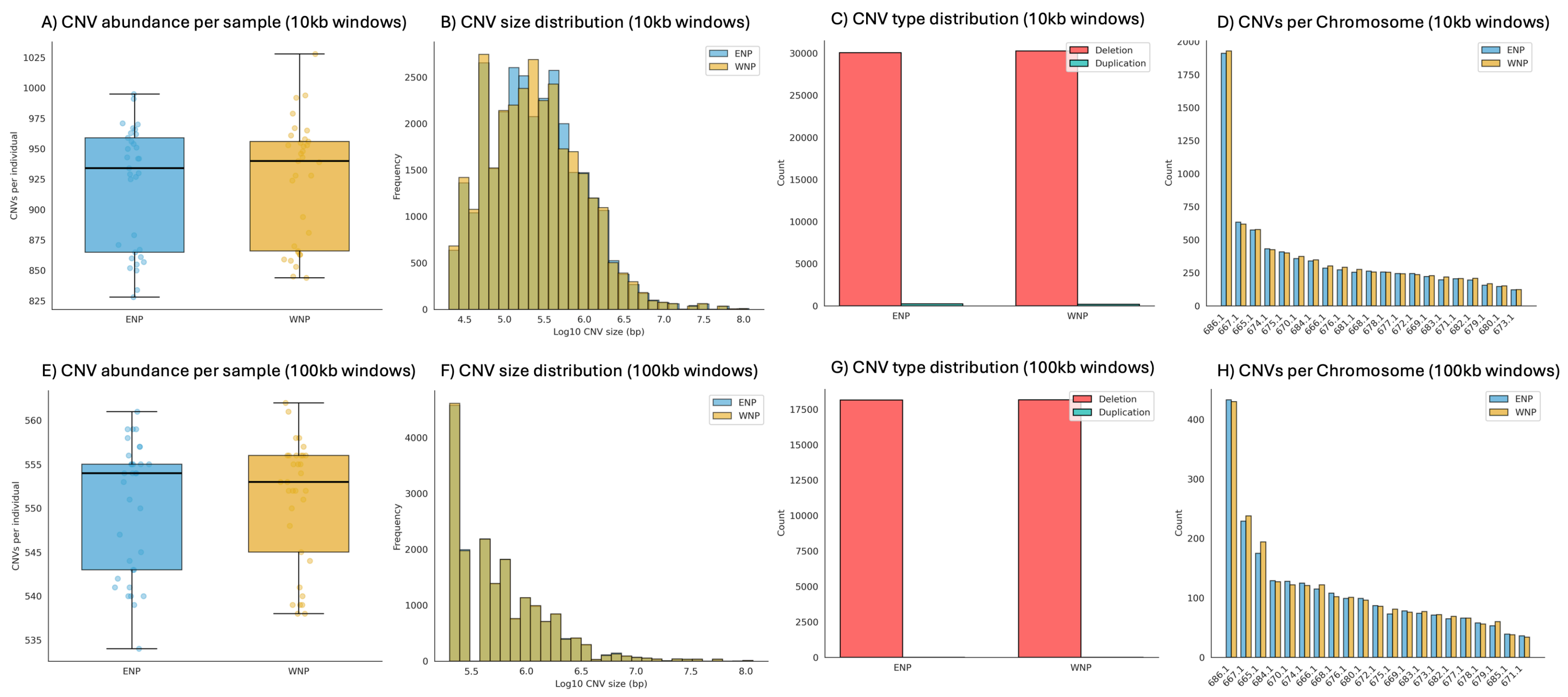


**Supplementary Figure 9:** Plots comparing copy number variants in whales sampled in the ENP and WNP. Panels A-D show CNV results for 10kb windows while panels E-H show CNV results for 100kb windows. A,E) CNV counts per sample. B,F) Distribution of CNV size in bp. C,G) Counts of duplications versus deletions. D,H) Counts of CNVs per chromosome.

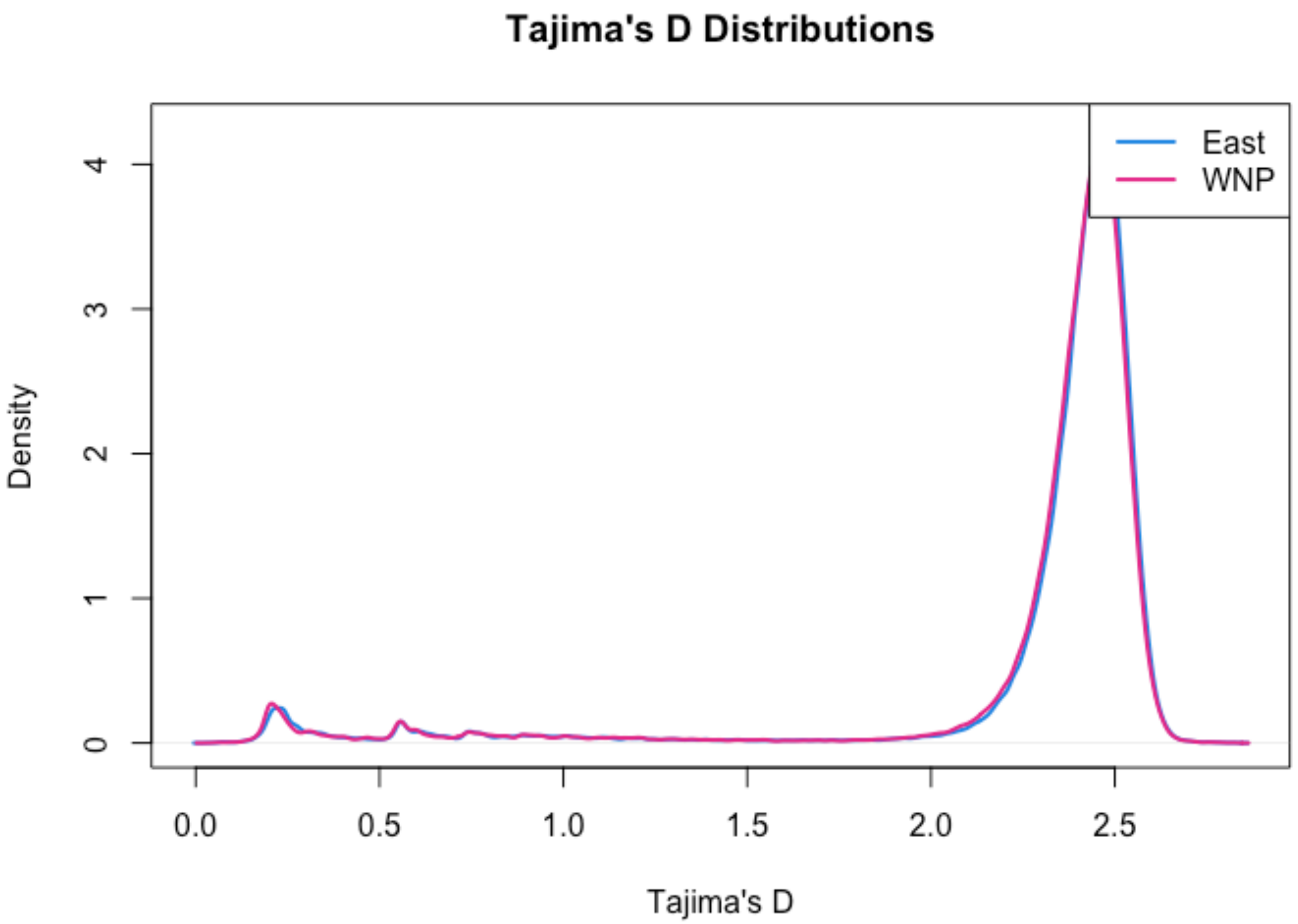


**Supplementary Figure 10:** Distribution of Tajima's D values across genomic windows for each sample site, where curves show the relative frequency of windows with each Tajima's D value.

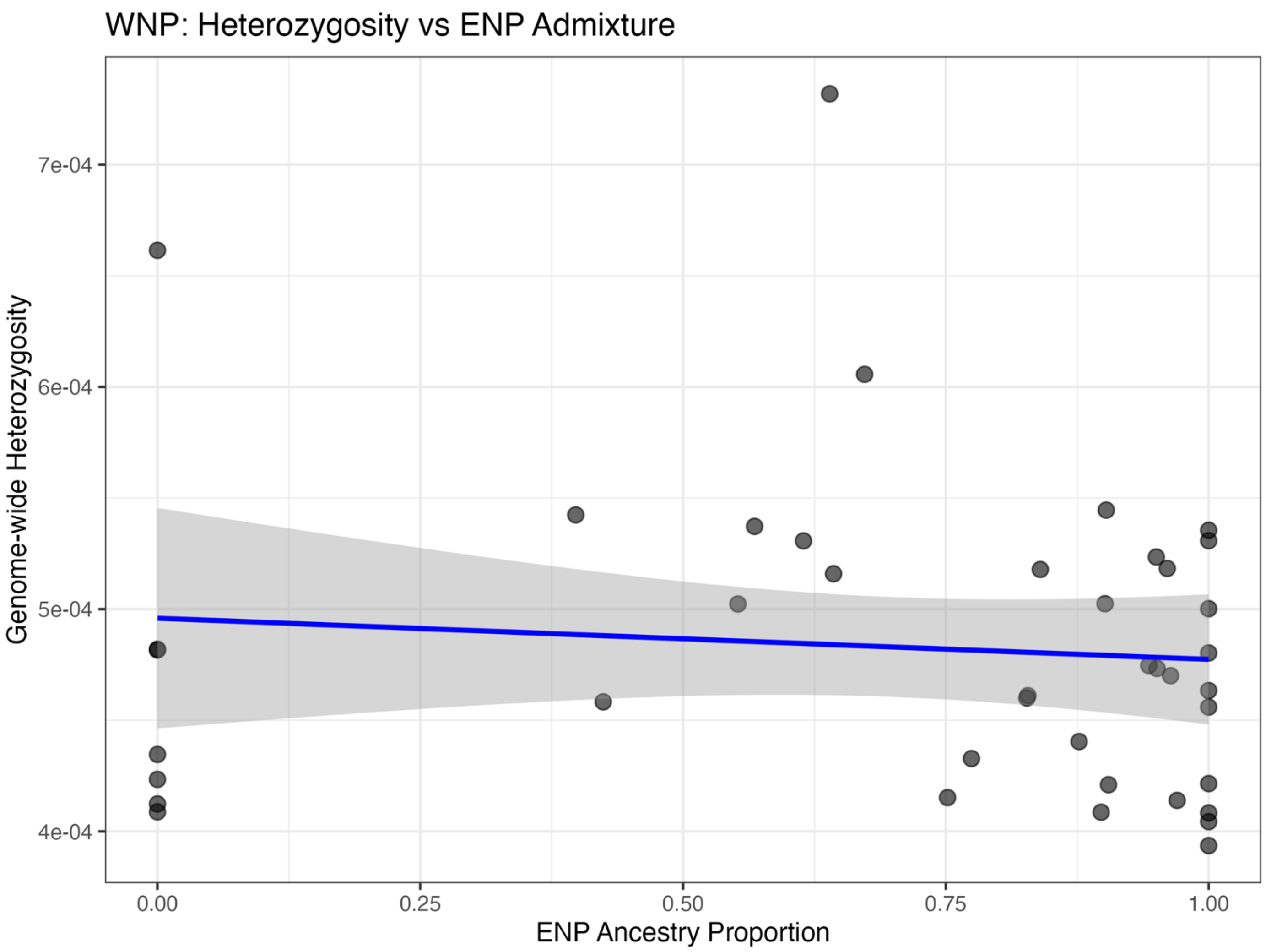


**Supplementary Figure 10:** Genome-wide heterozygosity plotted against admixture coefficients in WNP individuals. We found no significant relationship between the two.

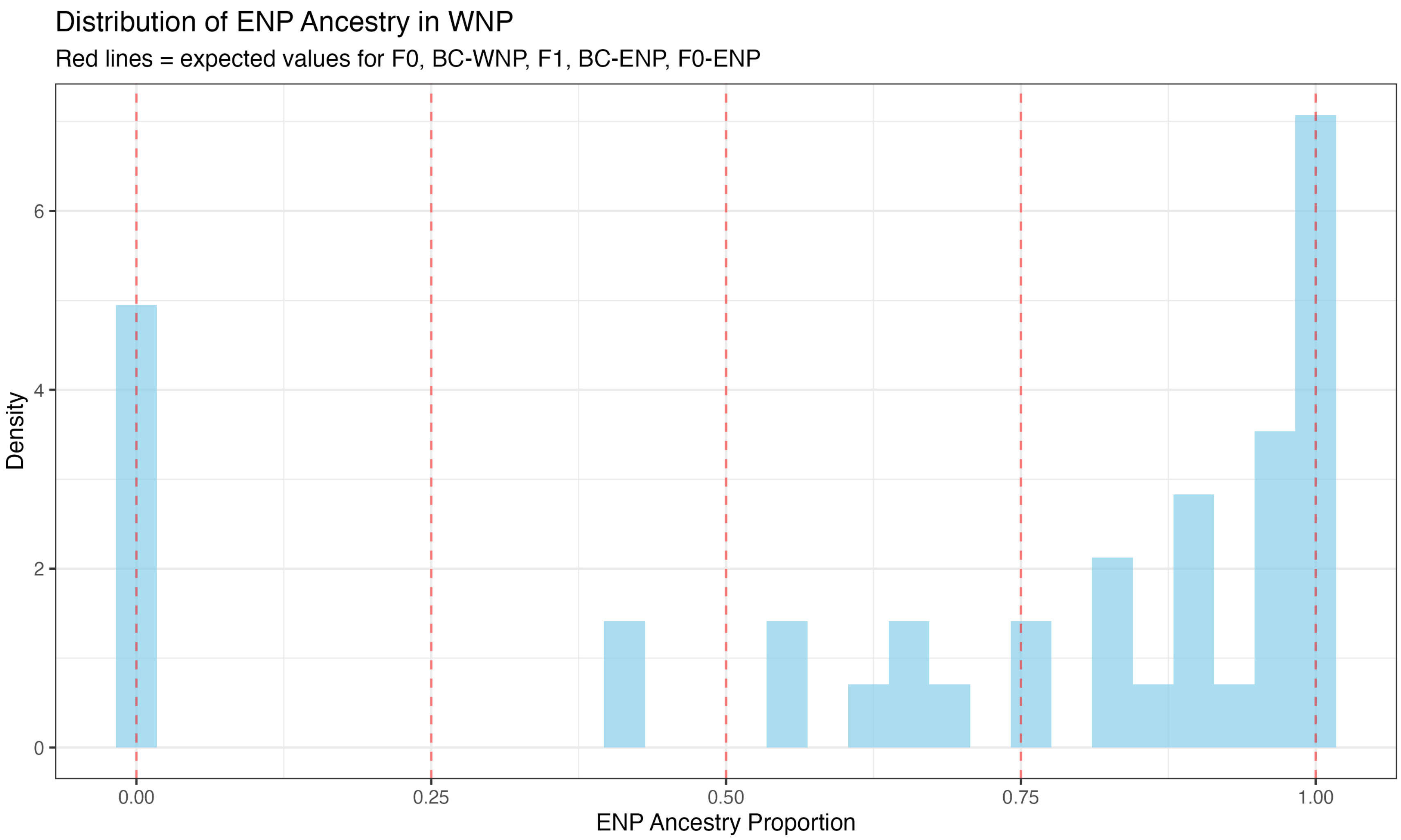


**Supplementary Figure 11:** Histogram of admixture coefficients in WNP individuals. We found no evidence of F1 hybrids or backcrosses based on admixture coefficient proportions.

**Supplementary Table 1:** Results of using the Evanno method to calculate ΔK for values of K from 1 to 5. K = 2 was the best supported value.

| K | Mean | Standard Deviation | ΔL | ΔK |
|---|---|---|---|---|
| 1 | -123373768 | 1.43E-06 | | 2.183E+12 |
| 2 | -120252105 | 1.77450711 | 3121663.08 | 1402292.53 |
| 3 | -117763727 | 84482.3029 | 2488378.07 | 23.6907482 |
| 4 | -115762278 | 137382.457 | 2001448.97 | 13.4597622 |
| 5 | -113913143 | 135268.16 | 1849135.2 | |

**Supplementary Table 2:** Genomic $F_{ST}$ randomly subsampled in 100 to 50,000 windows. Mean $F_{ST}$ was consistent across window count, indicating the value is robust to sampling variance across the genome.

| Windows Sampled | Mean $F_{ST}$ | SD across resamples | 95% CI |
|---|---|---|---|
| 100 | 0.01920 | 0.00104 | [0.01724, 0.02130] |
| 1,000 | 0.01913 | 0.00034 | [0.01850, 0.01978] |
| 10,000 | 0.01914 | 0.00011 | [0.01893, 0.01934] |
| 50,000 | 0.01914 | 0.00004 | [0.01905, 0.01923] |
